# Shear flow as a tool to distinguish microscopic activities of molecular machines in a chromatin loop

**DOI:** 10.1101/2024.01.23.576811

**Authors:** Sandeep Kumar, Ranjith Padinhateeri, Snigdha Thakur

## Abstract

Several types of molecular machines move along biopolymers like chromatin. However, the details about the microscopic activity of these machines and how to distinguish their modes of action are not well understood. We propose that the activity of such machines can be classified by studying looped chromatin under shear flow. Our simulations show that a chromatin-like polymer with two types of activities (constant or local curvature-dependent tangential forces) exhibits very different behavior under shear flow. We show that one can distinguish both activities by measuring the nature of a globule-to-extended coil transition, tank treading, and tumbling dynamics.

---

Spatial organization and biological functions of polymers, like DNA, chromatin, actin, etc., are made possible by various molecular motors/enzymes operating on them [1–4]. The activity of such ATP-dependent molecular machines influences the 3D conformations of DNA/chromatin. Motors on biopolymers perform a diverse set of functions such as reading the genetic code (RNA polymerase), re-organize proteins along DNA (SWI/SNF, CHD, ISWI, etc.), and move along the track transporting appropriate cargo (Kinesin, Dynein, Cohesin, Myosin) [2]. While performing these tasks, they apply a local force on the track that may bend or distort the polymer [5–8]. The microscopic details of how each motor acts on the filaments can differ. Some of the motors act like force-dipoles tangential to the chain [9–11] while others translocate along the backbone [12, 13]. The simultaneous action of such multiple ATP-consuming motors on a biopolymer creates a fascinating active system. For example, loop extruding motors fold chromatin, motors in the SWI/SNF, ISWI family slide nucleosomes leading to activation or repression of genes [14–16], correlated active motion on the chromatin leads to compact domain formation [17, 18] and spontaneous segregation in chromosome [11, 19].

The precise details of the microscopic action of these motors on the track are poorly understood. Experiments like X-ray crystallography and cryo-electron microscopy typically measure the static/frozen structures and are unable to reveal the dynamics of the motors [20, 21]. Light microscopy experiments, on the other hand, do not have sufficient resolution to probe the action of nanometer-sized motors [7, 22]. Hence, it will be appropriate to have a macroscopic experimental method combined with physical simulations that can give insight into the microscopic action of these motors on the filament. Shear flow experiments have been used extensively in the past to gain a microscopic understanding of the conformational properties and fluctuations of passive polymers [23–29]. These experiments show that the chain’s local structural properties can influence the polymer’s macroscopic behaviour under shear. However, very little is known about how active polymers behave under shear, given that the underlying motors can have different modes of action.

In this letter, we show that active polymer, even with slight differences in microscopic activity, can exhibit distinct collective macroscopic behavior under shear flow. Hence, we propose that studying active ring polymer under shear can help us distinguish between the different modes of microscopic activity of motors. To demonstrate this, we present a model with two types of motors having different tangential activities and show that they behave very differently under shear. We show that, under shear, the polymer undergoes a discontinuous transition from a collapsed to an open state in the presence of one type of activity, while the other type of activity results in a continuous transition. Further, we demonstrate that comparing macroscopic measurements of tumbling and tank-treading frequencies of the polymer can distinguish the two microscopic activities.

## MODEL AND METHODS

We consider a ring polymer of chromatin made of 200 beads (N=200) with neighbouring beads connected by harmonic spring potential and non-neighbouring beads interacting via a purely repulsive WCA potential [30]. Two sets of such ring polymers are simulated, one with type-I activity and the other with type-II activity, and both are subjected to a simple shear flow. The schematic diagram in Fig. 1 depicts the ring with two activities under shear. In type-I activity, every bead experiences a tangential force that depends solely on the local unit tangent vector. Hence, it has a constant magnitude of the active force on each monomer irrespective of the local shape along the polymer contour [31], given by

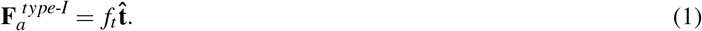

where,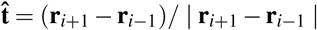 is the unit vector along tangent for any monomer *i* with position vector **r**_*i*_. On the other hand, in type-II activity, every bead experiences a tangential force that depends on the length of the tangent vector [32] given by

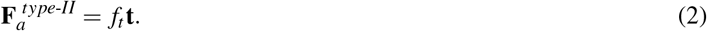

where, **t**_*i*_ = (**r**_*i*+1_ − **r**_*i*−1_)*/*2*l, l* =|**r**_*i*+1_ − **r**_*i*_| being the bond length. This implies that, unlike type-I, in type-II the local curvature along the contour affects the magnitude of active force. Each of the active r g polymers is subjected to a simple shear flow along the positive *x*-direction, with the velocity gradient tensor given by: 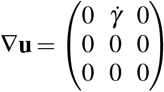, where the constant shear rate 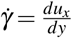.

**FIG. 1:**
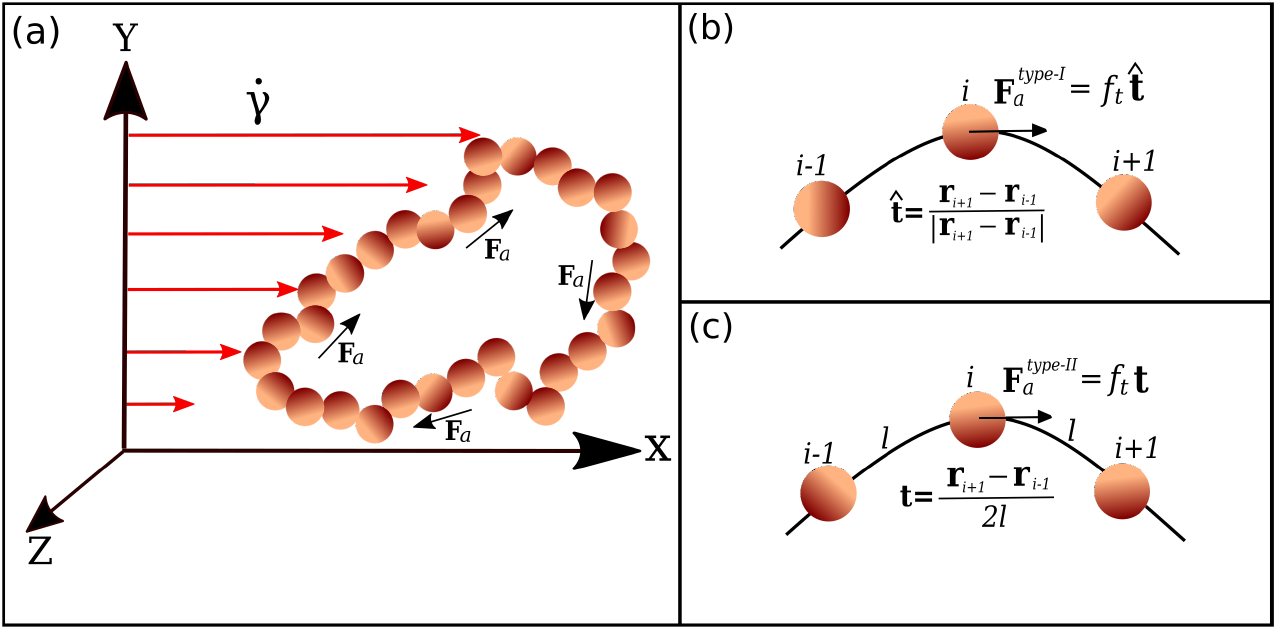
(a) Schematic representation of the active ring under the shear flow (with strength 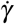). The flow is in the x-direction and the gradient is in the y-direction. (b) Type-I activity: constant magnitude of the active force 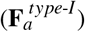. (c) Type-II activity: magnitude of the active force 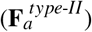 depends on local curvature. See text for details.

Fig. 1 shows the schematic for the active ring under shear. The equation of motion of the *i*^th^ bead is given by:

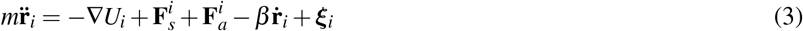

where *U*_*i*_ is the total interaction potential between the beads [30], **F**_**s**_ = *β* ∇**u** · **r** is the shear force with *β* being the friction coefficient, and ***ξ*** is the white noise [30].

The strength of the activity is measured by 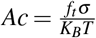 and we use the terms *Ac*-*I* (type-I) and *Ac*-*II* (type-II) to distinguish the two activities. The standard practice to represent the strength of the shear is to use the dimensionless Weissenberg number 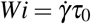, where *τ*_0_ is the longest relaxation time of the passive ring without shear. To determine *τ*_0_, we initialize the ring to an extended configuration with fractional extension *≈* 0.5. The ring is then allowed to relax without shear and active force. The longest relaxation time *τ*_0_ is then determined by fitting the decay of fractional extension to a single exponential *Aexp*(*−t/τ*_0_) + *B*. More details of the model can be found in supplementary material [30].

## RESULTS AND DISCUSSION

We quantify the structure of the ring by calculating the gyration tensor 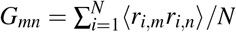, where *m, n ∈ {x, y, z}* and *r*_*i,m*_ is the position of *i*^*th*^ monomer in the centre of mass reference frame. The trace of *G*, 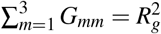, where *R*_*g*_ is the gyration radius of the polymer. Fig. 2 shows the heat map of *R*_*g*_ by varying *Ac* and *Wi*.

**FIG. 2:**
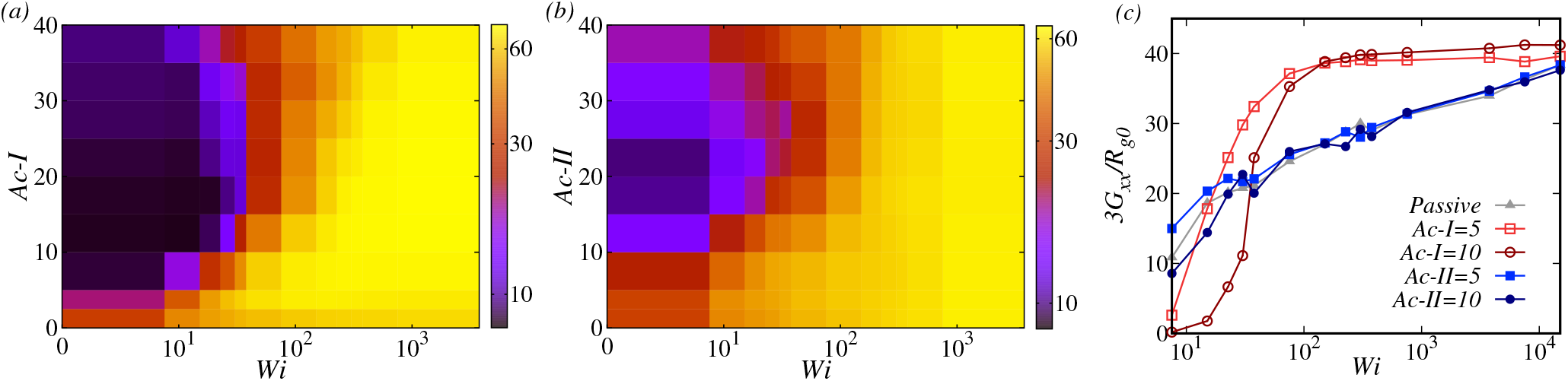
Heat map showing gyration radius *R*_*g*_ of the ring for different activity strength (*Ac*) and shear strength (*Wi*): (a) Activity type-I (*Ac*-*I*) and (b) Activity type-II (*Ac*-*II*). Type-I activity leads to a more compact configurations than type-II for lower *Wi*. (c) The component of the gyration tensor *G*_*xx*_ as a function of *Wi*.

### Without shear

The passive ring without shear is in the coiled state [33]. However, on increasing the activity in the absence of shear, we observe a coil-to-globule transition for both types of activities with subtle differences (see along the y-axis in Fig. 2 (a), (b)). One observes the transition to globular state for a lower value of *Ac*-*I* compared to *Ac*-*II* (See Fig. S2a). The collapse of the polymer with activity can be attributed to the emergence of local hairpin structures by the tangential force as characterized by the bond correlation (Fig. S2b). Further, the time for type-I to achieve the collapsed state is lower than type-II (Fig. S2c).

### Active and passive polymer with shear

The coiled passive ring starts to stretch on increasing *Wi* (see along the x-axis in Fig. 2 (a), (b)). The increase in *R*_*g*_ is by virtue of the monotonic increase in *G*_*xx*_ that squeezes the ring along the gradient (*y*) and vorticity (*z*) direction. The scaling of the components of gyration tensor *G*_*yy*_, *G*_*zz*_ *∼ Wi*^*−*0.45^ (Fig. S3) matches well with previous studies [34, 35]. The activity on the ring drives it to be in the globular state, whereas the shear tries to open it up. Hence, there is a stark difference between active and passive cases in the way the compactness of the polymer varies with *Wi* (compare *Ac* = 0 with *Ac >>* 0 in Fig. 2 (a),(b)).

### Rings with different activity types unfold differently under shear

We observe that the details of the structural transition under shear depend on the type of activity (*Ac*-*I* or *Ac*-*II*). As evident from Fig. 2, *Ac*-*I* supports more compact structure upto a larger *Wi* than *Ac*-*II*. Further, the *G*_*xx*_ shows a sudden increase in type-I compared to a gradual increase in type-II (Fig. 2(c)) as a function of *Wi*. Also, a substantial difference is observed in *G*_*zz*_, indicating more swelling in the vorticity direction for type-II (Fig. S3(c)).

Simulations give us the time series of the *R*_*g*_ (Fig. S4 and Fig. S5). From this, we compute the steady-state probability distribution of *R*_*g*_ and plot it in Fig. 3. The results for the passive case indicate the existence of the open state (a single predominant peak at large *R*_*g*_) for all *Wi* (Fig. 3(a)). However, comparison of passive *P*(*R*_*g*_) (Fig. 3(a)) with either of the active cases (Fig. 3(b),(c)) shows a clear difference between the two. For the active case, the *P*(*R*_*g*_) has a single closed state peak at low *Wi* followed by the double peak at intermediate *Wi*. The double peak is due to the competition between the activity and shear, resulting in switching between the open and closed states. At this intermediate *Wi*, we observe the following differences if we compare both activity types. A distinct bimodality in *P*(*R*_*g*_) is observed for *Ac*-*I*, showing a microscopic toggling between the open and closed state at intermediate *Wi*, which then finally goes to the open state. Note that the probability is zero for intermediate *R*_*g*_ values (between 30 to 50). However, the type-II activity always possesses finite weight for the intermediate *R*_*g*_s, indicating a gradual transition between closed to open state (also see Fig. 2(c)). It is this difference — the difference between the microscopic toggling and long-lived switching — is an important feature of our model that helps us differentiate between the two activities.

**FIG. 3:**
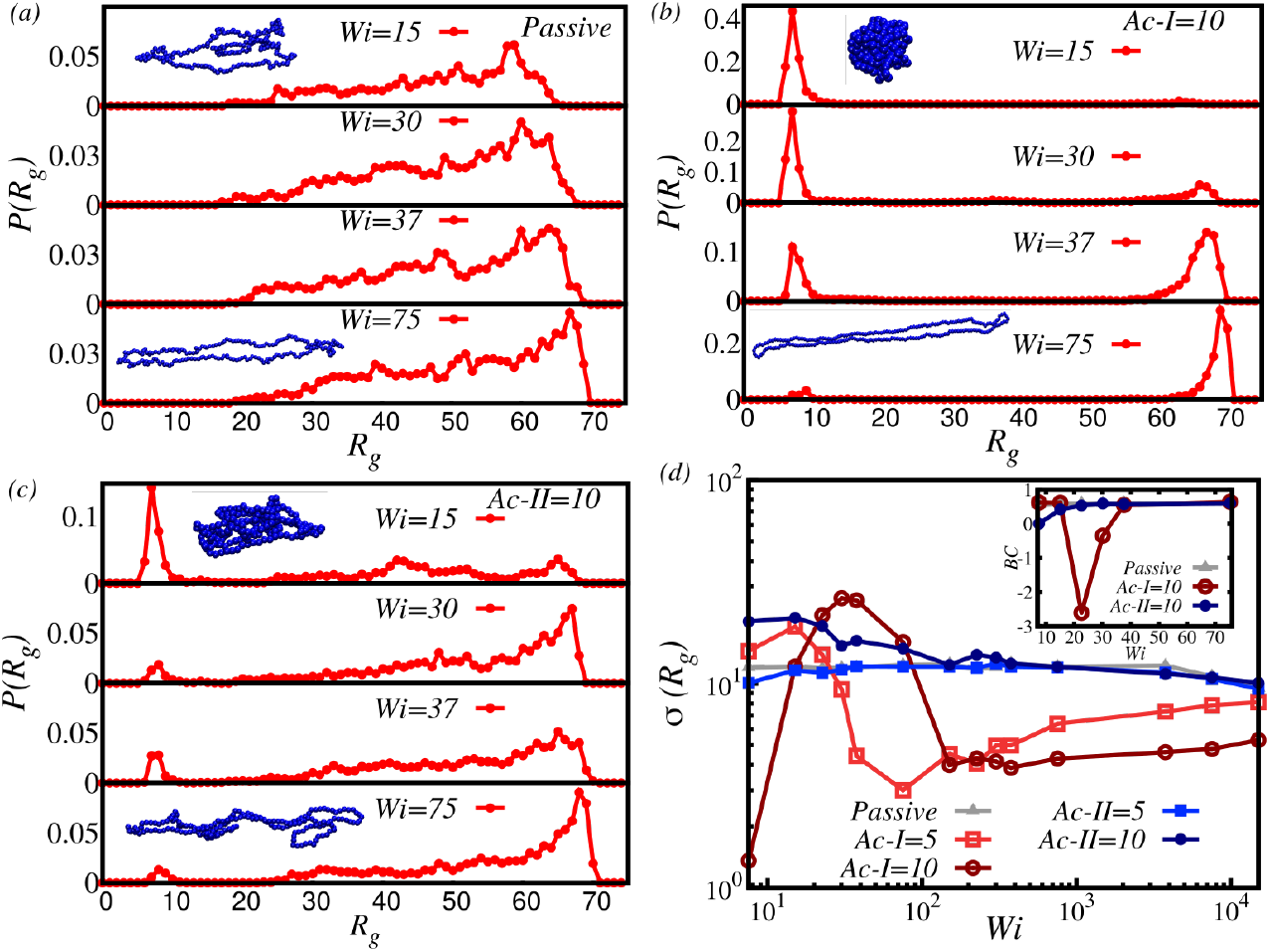
Distribution of *R*_*g*_ for (a) Passive, (b) *Ac*-*I* = 10 and (c) *Ac*-*II* = 10. The double peak in (b) and (c) shows the transition of the ring polymer from a collapsed state to a stretched state. Representative configurations of the polymer are shown in blue color. (d) Standard deviation and Binder cumulant of *R*_*g*_ suggest that transition under two types of activities are different.

To further quantify the difference between the two, we plot the standard deviation (*σ* (*R*_*g*_)) in Fig. 3(d). It is clear that passive, *Ac*-*I* and *Ac*-*II* curves are different. For *Ac*-*I*, the *σ* (*R*_*g*_) peaks at an intermediate *Wi* showing the signature of a discontinuous transition. On the other hand, *σ* (*R*_*g*_) for passive case is nearly flat while *Ac*-*II* is slightly higher than passive for low *Wi*. One can also see a clear difference between *Ac*-*I* and *Ac*-*II* at large *Wi*, indicating less fluctuations in the open state of type-I active ring. Further, the calculation of Binder cumulant [36–38], a well-known quantity to characterize the type of phase transition, supports the discontinuous nature of transition in type-I activity. The observed differences between the two activities under shear flow can help us experimentally identify the microscopic action of the active motors.

### Unlike tumbling (TB), tank-treading (TT) is more efficient in differentiating the activities

Polymers under shear are known to exhibit tumbling and tank-treading motion [26, 39–41]. Tumbling is the process where the entire polymer rotates about its centre of mass in the flow-gradient plane (See Movie 1 in [30]). This results from the net torque experienced by the polymer due to its extension along the gradient direction. The snapshots of the tumbling polymer are presented in Fig. S8. To quantify the tumbling motion, a cross-correlation (*C*_*xy*_) between conformational changes along flow and gradient direction is a standard mathematical tool [25, 42], defined as:

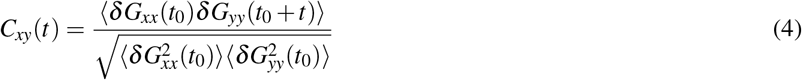

where *δ G*_*αα*_ = *G*_*αα*_ *− ⟨G*_*αα*_ *⟩* and 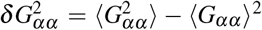 and the angular brackets represent the ensemble average over different *t*_0_. For a periodic tumbling motion, one would expect the *C*_*xy*_ to have an oscillatory behavior whose periodicity would define the tumbling frequency. The inset of Fig. 4 shows the *C*_*xy*_ for the two types of activities. The characteristic time of tumbling is obtained by *τ*_*tb*_ = 2(*t*_+_ *−t*_*−*_), where the times *t*_+_ *>* 0 corresponds to the deepest minimum and *t*_*−*_ *<* 0 for highest peak in *C*_*xy*_. In Fig. 4, we plot the scaled tumbling frequency *f*_*tb*_ = *τ*_0_*/τ*_*tb*_ for the passive and active rings. It is interesting to note that *f*_*tb*_ shows a power-law scaling as *f*_*tb*_ *∼ Wi*^0.68^ for both active and passive cases at higher *Wi*. For the passive case, such scaling has been observed before [25–27, 35, 39, 43–46]. The same scaling behavior for both types of activities at high *Wi* indicates the dominance of shear in this regime. Meanwhile, for the lower *Wi* regime, the two activities differ in two aspects. First, for the same activity strength, type-II exhibits tumbling at a smaller *Wi*. Secondly, the smoothness of tumbling is superior for type-II activity (see definition and details in [30]). Therefore, even though the frequency (*f*_*tb*_) is weak in differentiating the two activities, the smoothness of tumbling can capture the difference nicely.

**FIG. 4:**
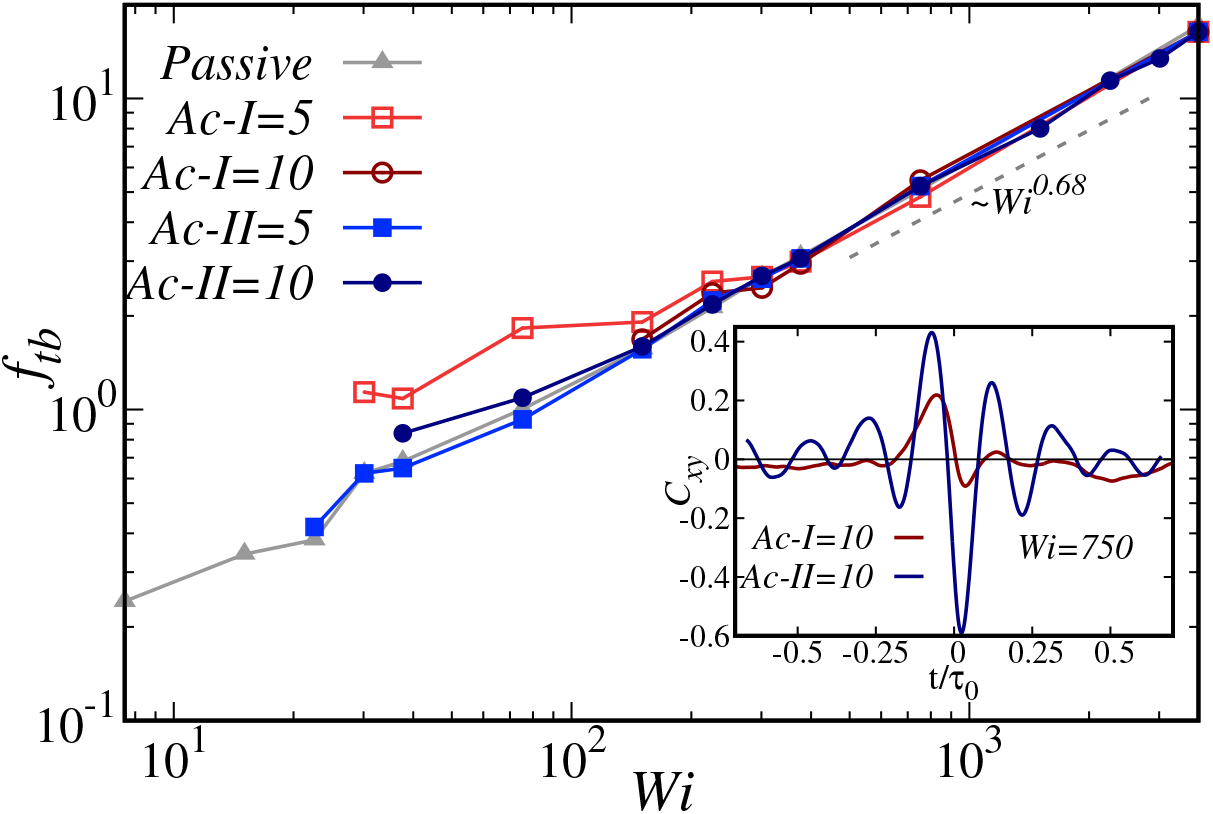
Scaled tumbling frequency *f*_*tb*_ for passive and active rings of both types at different *Wi* values showing the same power law for all cases at large *Wi*. For lower *Wi* the distinction between the two activities appear. Inset: Cross-correlation from the conformational fluctuations along the flow and gradient direction *C*_*xy*_ for *Ac*-*I, Ac*-*II* = 10. Note the difference between the activities.

In non-equilibrium situations like the shear flow, soft objects like vesicles and ring polymers are known to undergo tank-treading motion [25, 47–49]. In this mode, the polymer is stretched in the flow-gradient plane, forming an elliptical shape with monomers rotating about the center-of-mass along the contour (See Movie 2 in [30]). To identify and characterize the TT, we track the motion of monomers in the center-of-mass frame. For a given bead *i*, we define the fractional projection *X*_*i*_ = *x*_*i*_*/L*, where *x*_*i*_ = (**r**_*i*_*−* **r**_*cm*_)_*x*_ is the x-position of *i*^*th*^ monomer and *L* is the distance between the extreme beads along the longitudinal axis of the ring as shown pictorially in Fig. S12. The time auto-correlation of the *X*_*i*_ that will enable us to calculate the TT frequency can be defined as:

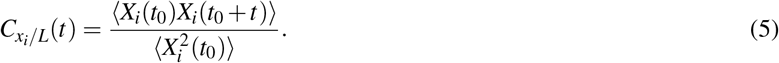

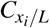 will exhibit an oscillatory motion for a perfect tank-treading. Fig. 5(a) shows distinct behaviors of 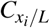 for the two types of activities. We obtain the characteristic time (*τ*_*tt*_) from the secondary peak position (the black dotted vertical line in Fig. 5(a)) and compute the tank-treading frequency as *f*_*tt*_ = *τ*_0_*/τ*_*tt*_. Fig. 5(b) plots the tank-treading frequency *f*_*tt*_ for passive and active rings. It is noteworthy that for passive rings, the *f*_*tt*_ *∼Wi*^0.72^, similar to the previous works [24, 25, 46, 50]. We find that even the active case approaches passive scaling at high *Wi*, suggesting the dominance of shear. At low *Wi*, both activities support TT, thereby enhancing the *f*_*tt*_ compared to its passive counterpart.

**FIG. 5:**
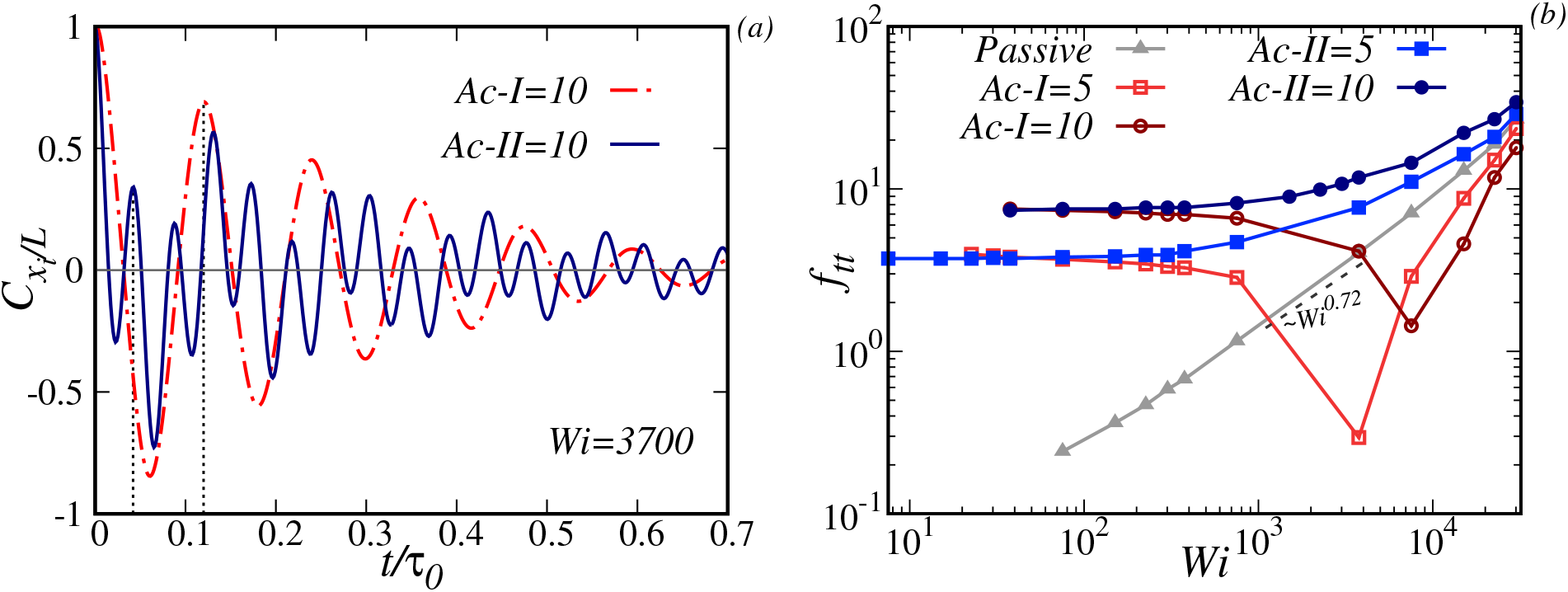
(a) The time auto-correlation of *X*_*i*_ (*C*_*x/L*_) for *Ac*-*I* = *Ac*-*II* = 10 and *Wi* = 3700. Note the difference in periodicity for the two activities. (b) Comparison of tank-treading frequencies (*f*_*tt*_) obtained from *C*_*x/L*_ for passive and both kinds of activities. The presence/absence of the dip differentiates both the activities.

However, at intermediate *Wi*, we find that both types of activities give very different TT features. This contrasting behaviour can differentiate the two activities. Around *Wi≈* 150, *f*_*tt*_ starts decreasing for *Ac*-*I*, it reaches a minimum and then rises until it approaches the passive curve. *Ac*-*II*, on the other hand, does not exhibit such non-monotonic behaviour. The minima in type-I activity correspond to the configurations where the ring polymer almost stops tank-treading and stalls (Fig S13 and Movie 3 in [30]). This is due to the interplay between the shear-induced TT and tangential force-induced TT. The shift in the minima implies that the stalling point is activity-dependent, suggesting that the shear required to balance it increases with activity. On further increasing *Wi*, shear overcomes the activity, and the ring tank treads only due to shear.

To gain more insights into the absence of minima in type-II, we re-examine the *R*_*g*_ distribution in Fig. 3. The distribution for type-II activity implies rings are more deformable in the open state (note more width around the prominent *R*_*g*_ peak) for type-II than type-I. A more flexible open state for type-II will help it to tumble; hence, there is no stalling, and the dynamics are a mix of TB and TT in type-II (Fig. S14 and Movie 4 in [30]).

To further show the distinction between the two types and examine the co-existence of TT and TB, we calculate the power spectral density (PSD) of *x*_*i*_*/L* time series (Fig. S13 and Fig. S14) by taking the Fourier transform of the time auto-correlation function *C*_*xi /L*_(*t*). The power spectrum 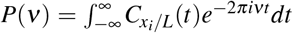 is plotted in Fig. 6 for both activities with two different shear strengths. Interestingly, type-I activity exhibits a single peak, whereas type-II shows a double peak in *P*(*ν*). The underlying feature of the single and double peaks can also be inferred from Fig. 5(a). The single peak in *Ac*-*I* is the evidence of long-lived TT dynamics. Whereas the double peak in *Ac*-*II* is the testimony of the co-existence of TB and TT dynamics, where one of the peaks corresponds to *f*_*tt*_ and the other is close to *f*_*tb*_.

**FIG. 6:**
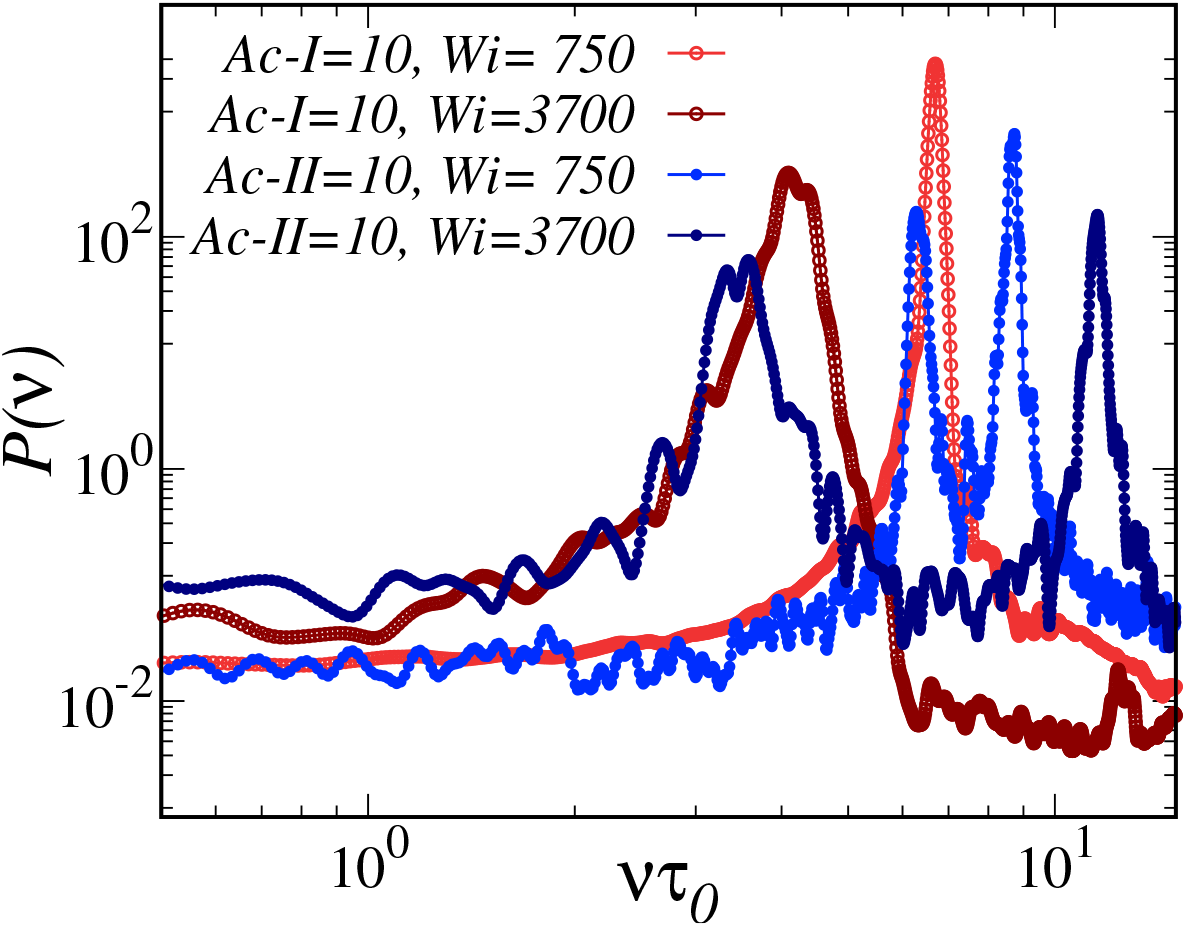
Power spectrum of *x/L* time-series for *Ac*-*I* = *Ac*-*II* = 10 for different *Wi*. Note that the *Ac*-*I* leads to distributions with a single peak, while the *Ac*-*II* contains a double peak in its distribution.

In conclusion, we have shown that one can employ shear as a tool to distinguish the dynamics of an active ring under the influence of two different types of microscopic activities. Our results show that the type of activity at the microscopic level strongly affects macroscopic features such as collapsed to open state transition under shear. In particular, the type-I activity exhibits a discontinuous collapse-to-open transition as opposed to a continuous transition for type-II. The dynamical responses of the active polymers under flow are also very different. For intermediate shears, type-II active rings support more tumbling, whereas type-I is more prone to tank treading. Such types of microscopic motor action are common in biological systems, particularly among motors that act on chromatin polymers. Hence, our simulations can be used to design future experiments on chromatin motors to classify their activity.

The authors would like to acknowledge the HPC facility at IISER Bhopal for all the computational work. ST acknowledges SERB India.

