## Supplementary Material for "Shear flow as a tool to distinguish microscopic activities of molecular machines in a chromatin loop"

#### S1 Model and Simulation Details

We consider an isolated flexible ring polymer of size $N = 200$, which is subjected to simple shear flow. Neighboring monomers are connected by harmonic spring potential, which is defined as :

$$U_h = \frac{1}{2}\kappa(r_{ij} - l_0)^2 \tag{1}$$

where, $\kappa$ is the spring constant and $l_0$ is the equilibrium separation between the monomers. The non-neighboring monomers interact via purely repulsive L-J potential (WCA) to prevent overlap between them.

$$U_{LJ} = \begin{cases} 4\epsilon\left[\left(\frac{\sigma}{r}\right)^{12} - \left(\frac{\sigma}{r}\right)^{6} + \frac{1}{4}\right] & \text{for } r = |\mathbf{r}_i - \mathbf{r}_j| < 2^{\frac{1}{6}}\sigma \\ 0 & otherwise \end{cases} \tag{2}$$

Here, $\epsilon$ is the strength of the repulsion and $\sigma$ is the diameter of each monomer.

To include activity on the polymer we apply two types of tangential force on the polymer. In type-I activity, a tangential force is applied using the following definition:

$$\mathbf{F}_a^{\textit{type-I}} = f_t\hat{\mathbf{t}}. \tag{3}$$

where $\hat{\mathbf{t}} = (\mathbf{r}_{i+1} - \mathbf{r}_{i-1})/ \mid \mathbf{r}_{i+1} - \mathbf{r}_{i-1} \mid$ is the unit vector along the tangent to the backbone of $i^{th}$ monomer. It is to be noted that in equation(3), i.e. in type-I activity, the magnitude of active force remains constant irrespective of the shape fluctuations of the polymer contour because the force is applied along the unit vector. In a separate set of simulations, we adopt another form of the active force defined as :

$$\mathbf{F}_a^{\textit{type-II}} = f_t\mathbf{t}. \tag{4}$$

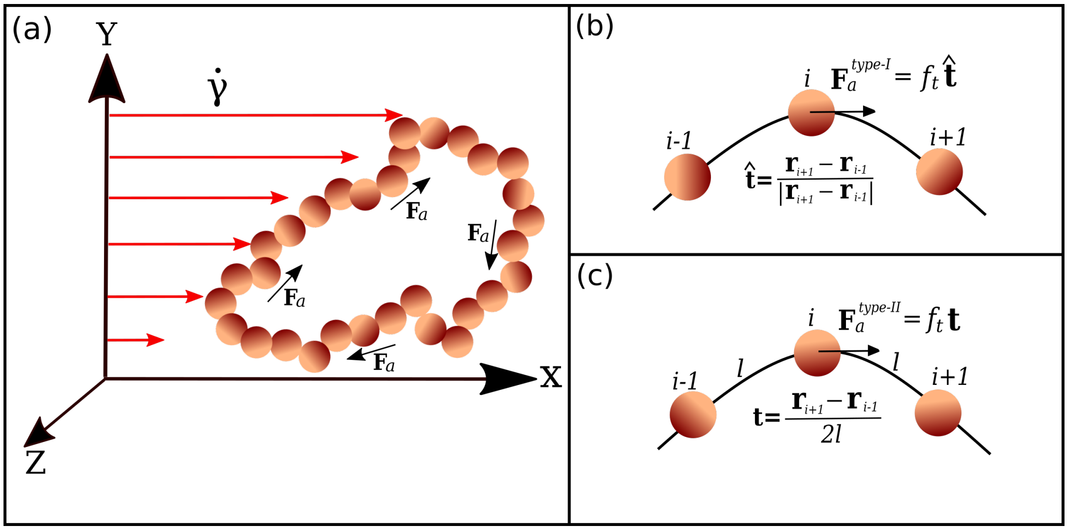


Figure S1: This is the reproduction of figure 1 of main text, showing the schematic representation of active ring under the shear flow.

$\mathbf{t}_i = (\mathbf{r}_{i+1} - \mathbf{r}_{i-1})/2l$, where $l =| \mathbf{r}_{i+1} - \mathbf{r}_i |$ is the bond length. Note that type-II activity constitutes the active force that is affected by the local curvature along the contour.

Fig. S1 shows the schematic representation of our active ring polymer subjected to simple shear flow along the x-direction. Because of the shear flow, the velocity field is modified by a velocity gradient tensor given by : $\nabla \mathbf{u} = \begin{pmatrix} 0 & \dot{\gamma} & 0 \\ 0 & 0 & 0 \\ 0 & 0 & 0 \end{pmatrix}$, $\dot{\gamma} = \frac{du_x}{dy}$ is the constant shear rate. Therefore, for each $i^{th}$ bead the equation of motion is given:

$$m\ddot{\mathbf{r}}_i = -\nabla U_i + \mathbf{F}_s^i + \mathbf{F}_a^i - \beta \dot{\mathbf{r}}_i + \boldsymbol{\xi}_i \tag{5}$$

$\mathbf{F_s} = \beta \nabla \mathbf{u} \cdot \mathbf{r}$ is the force due to shear flow, $\mathbf{F}_a$ is the active force, $\beta$ is the friction coefficient and $\boldsymbol{\xi}$ is the Gaussian white noise with zero mean and unit variance and satisfies the relation $\langle \boldsymbol{\xi}_i(t_1) \cdot \boldsymbol{\xi}_j(t_2) \rangle = \sqrt{6\beta k_B T} \delta_{ij} \delta(t_1 - t_2)$.

The strength of the activity is measured by the activity number $Ac = \frac{f_t \sigma}{K_B T}$. To distinguish between the two forms of active force, we use $Ac$-$I$ and $Ac$-$II$ to measure the activity strength for type-I and type-II rings, respectively. Apart from that, we use the well-known quantity called Weissenberg number ($Wi$) to account for the strength of the shear flow. $Wi = \dot{\gamma}\tau_0$, where $\tau_0$ is the longest relaxation time of the passive ring without shear. To calculate $\tau_0$, we start from the maximum extended configuration of the ring polymer in the x-direction, for which the fractional extension $\Delta x/L \approx 0.5$, and let the system evolve without shear and active force. The ratio $\Delta x/L$ decays with time until it reaches saturation. We fit the tail end of the decay (when $\Delta x/L \leq 0.2$) with the expression $(\Delta x/L)^2 = Aexp(-t/\tau_0) + B$ to obtain the longest relaxation time $\tau_0 \approx 7500$.

*Simulation Parameters:* In the present study, the ring polymer is placed near the centre of a cubic box of size $200 \times 200 \times 200$. All simulations are performed at a constant temperature of the thermal bath with $K_BT = 0.1$ and $\epsilon = 0.1$. The equilibrium separation between monomers is $l_0 = 1.25\sigma$, where $\sigma = 2$ is the diameter of each bead. The spring constant $\kappa$ has been taken to be $250, 1000, 3000$ depending on the shear rate to keep $l_0$ fixed. To evolve the system, we integrate the equation of motion using the velocity-Verlet scheme in steps of $dt = 10^{-3}$ and $dt = 5 \times 10^{-4}$. The polymer is allowed to relax under shear flow for more than $10 \times \tau_0$, in more than 40 independent sets of simulations. Shear is applied to the system using Lees-Edwards boundary conditions.

### S2 Structural analysis

In Fig. S2(a), we show the effect of type-I and type-II active force on polymer size in the absence of shear. Without shear, both kinds of activity eventually lead to the collapsed state of the ring. However, a higher magnitude of activity is required to trigger the collapse in type-II ($Ac$-$II = 10$ onwards) than type-I ($Ac$-$I = 5$ onwards). Once the globular state is reached, the gyration radius remains similar in both cases (Fig. S2(a)). As shown in our previous study [1], the collapse of the ring is due to the formation of local hairpin-like structures. In Fig. S2(b), we plot the bond-correlation function defined as $\beta(s) = \langle \mathbf{b}_{i+s} \cdot \mathbf{b}_i \rangle$, where $\mathbf{b}_i = \mathbf{r}_{i+1} - \mathbf{r}_i$ is the bond vector. The negative minimum in $\beta(s)$ is a signature of such local hairpin structures, which are present in $Ac$-$I \geq 5$ and $Ac$-$II \geq 10$. Further, it is also observed in Fig. S2(c) that $\mathbf{F}_a^{type-II}$ takes a longer time to collapse the ring as compared to $\mathbf{F}_a^{type-I}$.

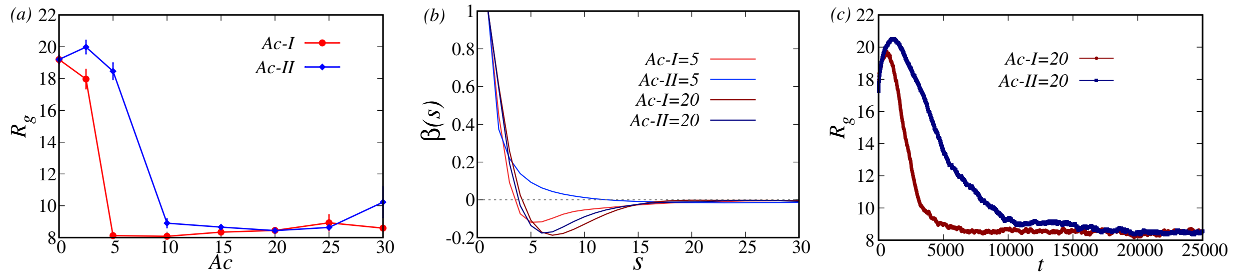


Figure S2: (a) Comparison of polymer compaction for the two kinds of active forces without shear ($Wi = 0$). (b) Bond correlation of type-I and type-II active ring without shear($Wi = 0$). (c) $R_g$ vs $t$ for $Wi = 0$ case showing faster collapse in type-I active ring.

The gyration tensor ($G_{mn}$) analysis of the ring in the presence of shear flow shows a considerably distinct effect of the kind of microscopic activity on the polymer structure. In Fig. S3, we show the scaled diagonal elements of the gyration tensor, $3G_{xx}/R_{g0}$, $3G_{yy}/R_{g0}$ and $3G_{zz}/R_{g0}$ where $R_{g0}$ is the square of gyration radius without shear. Passive ring polymers show a steady rise in $G_{xx}$ with increasing $Wi$ as they get stretched in the flow direction. This elongation in the x-direction occurs at the expense of the polymer's compaction along the gradient (y) and vorticity (z) direction, as seen in Fig. S3(b) and (c). The type-II ring displays a similar behaviour of $G_{mm}$ as that of passive with a small deviation for lower $Wi$. However, type-I rings show a sudden transition from globular to completely extended state with $G_{xx}$

saturating after $Wi \approx 150$. In the gradient direction, $G_{yy}$ also displays a small region of non-monotonicity at low $Wi$. In this region, the globular ring polymer unfolds steadily, thereby increasing $G_{yy}$ and reaching a maximum value. Further increasing $Wi$ leads to compaction along the y-direction, and $G_{yy}$ decreases as $G_{xx}$ increases rapidly. Interestingly, we observe a swelling of type-II ring polymer in the vorticity direction compared to type-I shown in the $G_{zz}$ plot.

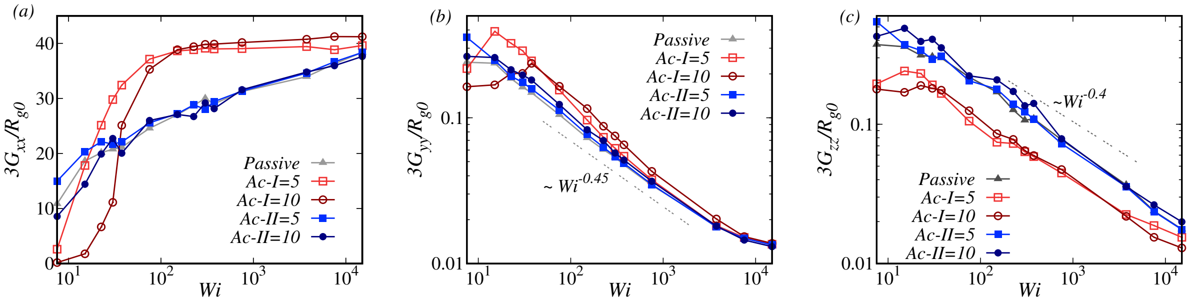


Figure S3: $G_{xx}, G_{yy}$ and $G_{zz}$ as a function of $Wi$ for both kind of tangential activity.

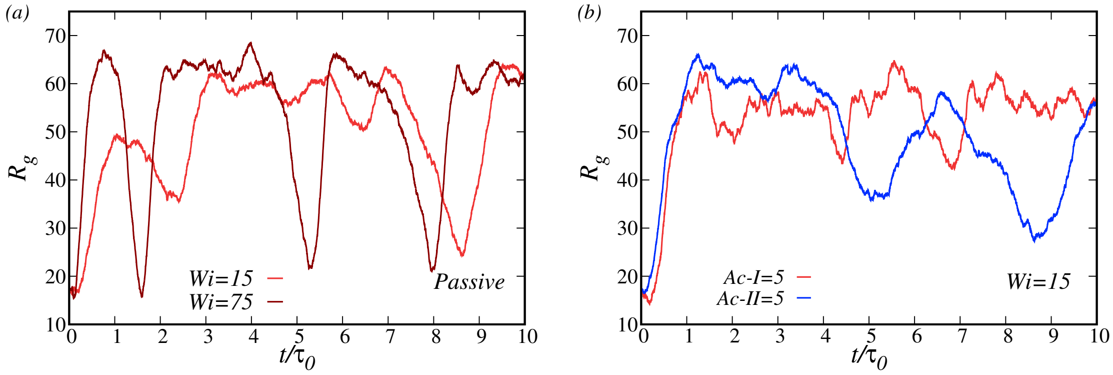


Figure S4: Time evolution of $R_g$ for passive and $Ac\text{-}I, Ac\text{-}II = 5$.

In Fig. S4 and Fig. S5, we present the time series of $R_g$, which has been used to calculate probability distribution ($P(R_g)$) in the main text as well as in Fig. S6 and Fig. S7. The passive ring in Fig. S4(a) spans a wide range of $R_g$ values, which leads to a broad distribution of $P(R_g)$. Distinct open and closed states for type-I activity are also visible here. For $Ac\text{-}I = 10$, the ring remains in the globular state at $Wi = 15$ and stretches completely at $Wi = 75$. It should be noted that type-I remains very stable in either collapsed or stretched configurations, which gives rise to a bimodal distribution of $R_g$. This feature is absent in type-II rings, which makes a smooth transition between compact and elongated states. In the subsequent figures (Fig. S6 and Fig. S7) we present the $P(R_g)$ for $Ac\text{-}I = 5, 20$ and $Ac\text{-}II = 5, 20$, which shows similar features as discussed above. We also calculate the standard deviation of the probability distribution $P(R_g)$ as $\sigma(R_g) = \sqrt{\sum_{i=1}^{n}(R_g^i - \langle R_g \rangle)^2/n}$,

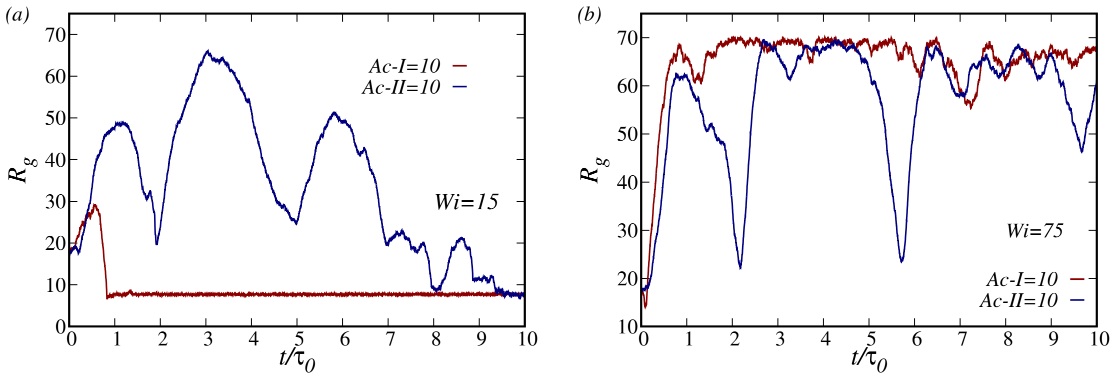


Figure S5: Time evolution of $R_g$ for *Ac-I*, *Ac-II* $= 10$ at $Wi = 15$ and 75.

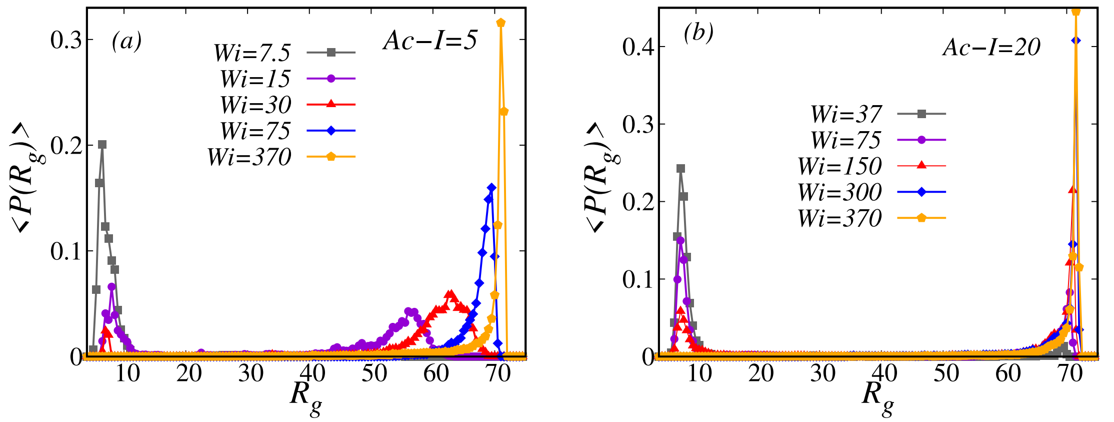


Figure S6: Distribution of $R_g$ for *Ac-I* $= 5$ and 20 at different $Wi$ values.

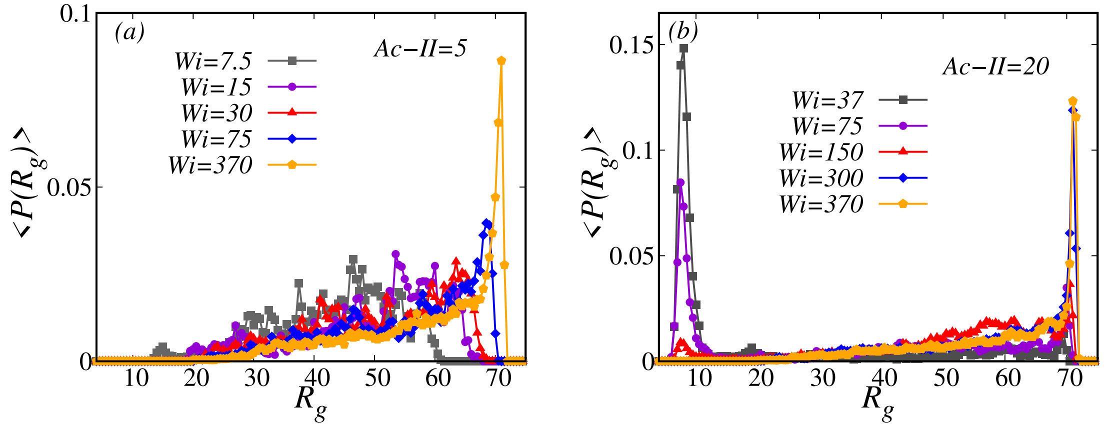


Figure S7: Distribution of $R_g$ for *Ac-II* $= 5$ and 20 at different $Wi$ values.

which is shown in the main text. The binder cumulant for $R_g$ distribution is defined as $BC = 1 - \frac{\langle R_g^4 \rangle}{3\langle R_g^2 \rangle^2}$ and has been discussed in the main text.

### S3 Tumbling Properties

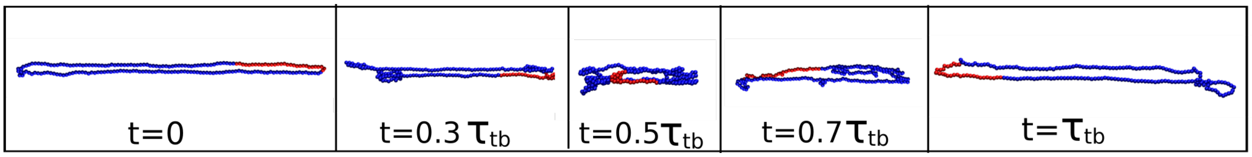


Figure S8: Snapshot of an active ring polymer showing tumbling motion at $Wi = 370$. Time is denoted in increasing order from $t = 0$ to $t = \tau_{tb}$. The red segment in the ring is for visual aid.

Snapshots of a complete tumbling event are shown in Fig. S8. The polymer undergoes one tumbling cycle in time interval $t = 0$ to $t = \tau_{tb}$, where $\tau_{tb}$ is the characteristic tumbling time. As discussed in the main text, the characteristic tumbling time $\tau_{tb}$ as well as tumbling frequency $f_{tb}$ can be obtained from the cross-correlation function given as:

$$C_{xy}(t) = \frac{\langle \delta G_{xx}(t_0)\delta G_{yy}(t_0 + t)\rangle}{\langle \delta G_{xx}^2(t_0)\rangle\langle \delta G_{yy}^2(t_0)\rangle} \tag{6}$$

where, $G_{\alpha\alpha}$ are elements of the gyration tensor and $\delta G_{\alpha\alpha} = G_{\alpha\alpha} - \langle G_{\alpha\alpha}\rangle$ and $\delta G_{\alpha\alpha}^2 = \langle G_{\alpha\alpha}^2\rangle - \langle G_{\alpha\alpha}\rangle^2$ . In Fig. S9, we report the $C_{xy}$(t) as a function of time for passive, $Ac\text{-}I = 10$

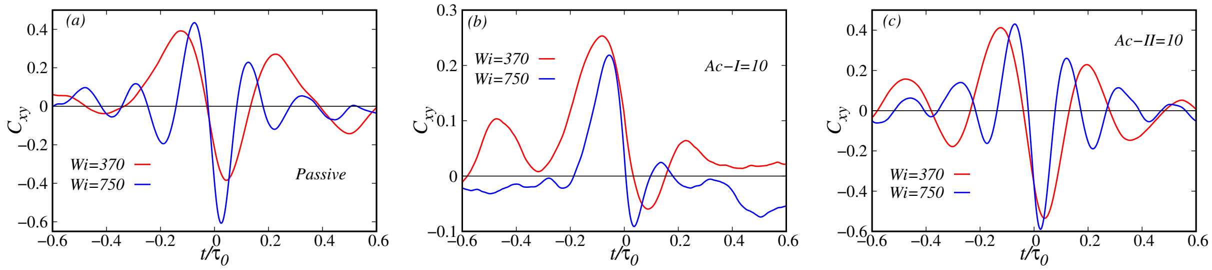


Figure S9: Cross-correlation ($C_{xy}$) of the fluctuations in $G_{xx}$ and $G_{yy}$ for passive, $Ac\text{-}I = 10$ and $Ac\text{-}II = 10$ case.

and $Ac\text{-}II = 10$, which is then utilized to calculate the tumbling frequency $f_{tb}$ as defined in the main text. As discussed there, $f_{tb}$ shows similar scaling in type-I and type-II rings; however, it is clear from Fig. S9 that the amplitudes of $C_{xy}$ in type-I rings are much smaller as compared to passive and type-II rings. The amplitude can be related to the smoothness of tumbling. We quantify the smoothness from the height of the positive and negative peaks and define it as $Q_{tb} = C_{xy}^{+} - C_{xy}^{-}$, where $C_{xy}^{+}$ and $C_{xy}^{-}$ are the heights of positive and negative peaks of $C_{xy}$ respectively. $Q_{tb}$ shown in Fig. S10 is observed to be higher in passive and type-II rings as compared to type-I rings. The $C_{xy}$ plot, along with their $Q_{tb}$ values, suggest that the tumbling events are less smooth in the case of type-I, i.e. the constant active force case. Here, the active force always tries to reduce the fluctuations in the polymer size along the gradient direction, which in turn makes $\delta G_{yy}$ small. Although this mechanism does not affect the tumbling frequency, rather it makes the tumbling events less smooth. On the other hand, when activity varies according to the local shape of the polymer backbone (type-II), those random fluctuations ($\delta G_{yy}$) are sustained, which consequently leads to very clear tumbling events.

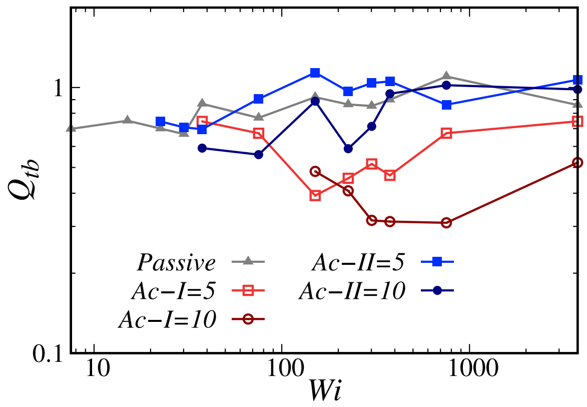


Figure S10: Smoothness of tumbling $Q_{tb}$ for the two case of tangential activity.

### S4 Tank-Treading Properties

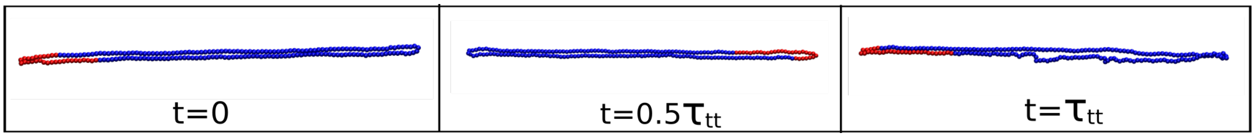


Figure S11: A representative snapshot of an active ring polymer showing Tank-treading motion. Time is denoted in increasing order from $t = 0$ to $t = \tau_{tt}$. The red segment in the ring is for visual aid.

Ring polymers subjected to shear flow show a peculiar motion known as tank-treading (TT), in which the ring maintains its shape and rotates steadily about its centre of mass. TT motion occurs when the polymer is completely stretched in the flow-gradient plane with a high velocity gradient across monomers. In Fig. S11, we show the snapshots of an active ring undergoing a complete tank-treading cycle. As evident from the snapshots, the segment coloured in red moves from the left side (at $t = 0$) to the right side (at $t = 0.5\tau_{tt}$) and again comes back to the left side after completing the complete TT cycle (at $t = \tau_{tt}$). To comprehend the characteristic time or frequency of the TT cycle, we track the motion of a monomer in the following way. For a given bead $i$, we define $X_i = x_i/L$, where $x_i = (\mathbf{r}_i - \mathbf{r}_{cm})_x$ is the x-position of $i^{th}$ monomer in the centre of mass ($r_{cm}$) reference frame and $L$ is the distance between the extreme beads along the longitudinal axis of the ring. A pictorial representation of $x_i$ and $L$ is given in Fig. S12.

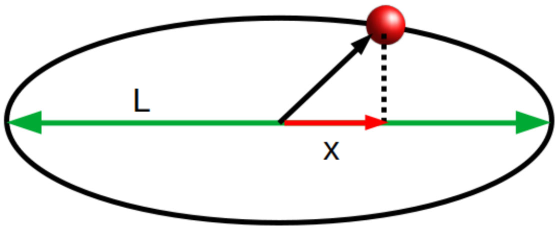


Figure S12: Pictorial representation of the longitudinal axis $L$ shown in green. $\mathbf{x}$ is the projection of an arbitrary monomer's position on the longitudinal axis.

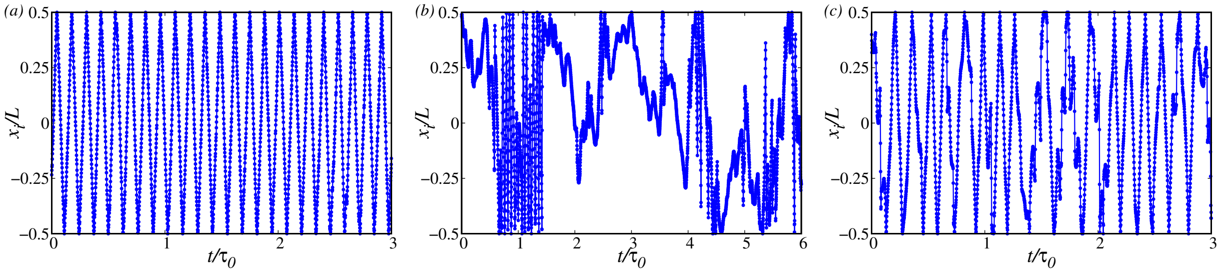


Figure S13: Variation of $x_i/L$ for $Ac$-$I = 5$ at $Wi = 370, 3700\ \&\ 7500$

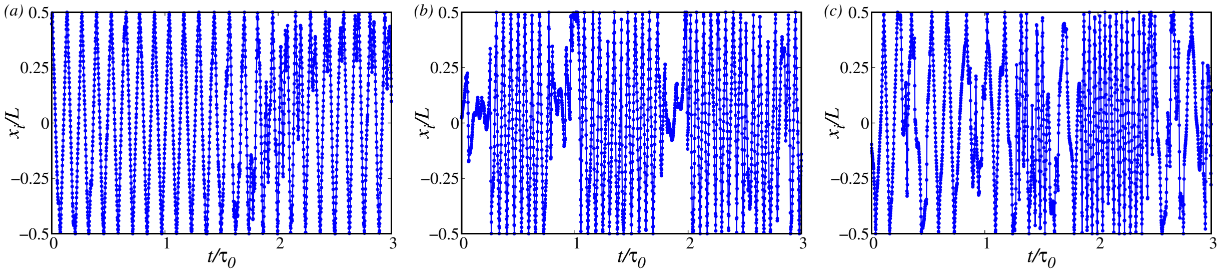


Figure S14: Variation of $x_i/L$ for $Ac$-$II = 5$ at $Wi = 370, 3700\ \&\ 7500$.

Fig. S13 shows the variation of $x_i/L$ for $Ac$-$I = 5$ at three different $Wi$ values. At $Wi = 370$, $X_i$ displays a nice oscillatory behaviour with time as the monomers move along the elliptical path on the contour during the TT cycle (Fig. S13(a)). However, as we increase $Wi$ to 3700 (Fig. S13(b)), competition between the constant active force and shear flow begins to appear, and the TT motion slows down. This phenomenon is evident from the irregular oscillation in $X_i$ in Fig. S13(b). Upon further increasing the $Wi$, shear dominates the activity, and TT motion is restored. In contrast, for $Ac$-$II = 5$ shown in Fig. S14, this competition between the active force and shear is hugely suppressed, and the ring exhibits TT motion at all values of $Wi$.

We define the time auto-correlation of the fraction $x_i/L$ as:

$$C_{x_i/L}(t) = \frac{\langle X_i(t_0) X_i(t_0 + t)\rangle}{\langle X_i^2(t_0)\rangle} \tag{7}$$

and plot it in Fig. S15.

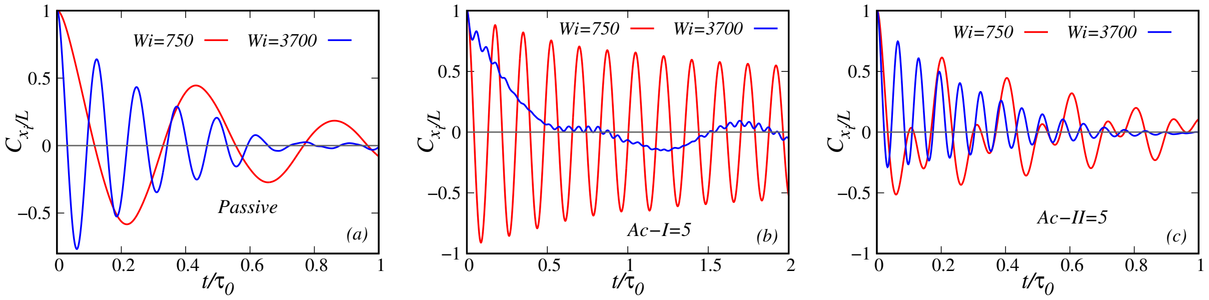


Figure S15: $C_{x/L}$ for passive, $Ac\text{-}I = 5$ and $Ac\text{-}II = 5$ case.

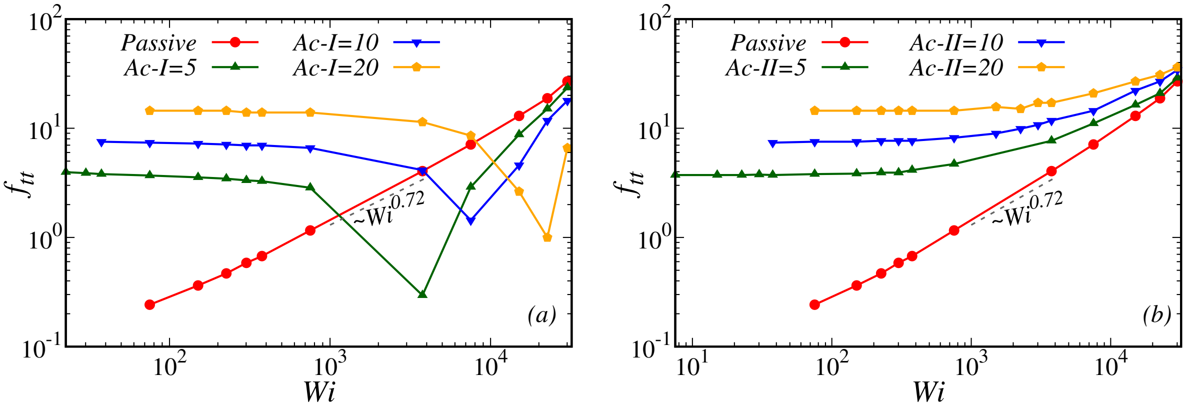


Figure S16: Scaled tank-treading frequency for type-I and type-II active rings at different $Wi$ obtained from $C_{x/L}$.

The secondary peak in the plot of $C_{x_i/L}(t)$ is used to identify $\tau_{tt}$, which is the characteristic time of tank-treading motion. We obtain scaled tank-treading frequency $f_{tt} = \tau_0/\tau_{tt}$. In Fig. S15, we report the $C_{x_i/L}$ for passive, $Ac\text{-}I = 5$ and $Ac\text{-}II = 5$ at two different $Wi$. The corresponding tank-treading frequency for type-I and type-II active rings has been shown separately in Fig. S16. For the passive ring, $f_{tt}$ increases monotonically with an increase in $Wi$ following a power law scaling of $\sim Wi^{0.72}$.
On the other hand, the active rings show a completely different response in TT frequency when $Wi$ is increased. It is noteworthy that, unlike the passive case, the active ring at lower $Wi$ exhibits a plateau in the tank-treading frequency $f_{tt}$ whose magnitude increases with activity strength. This is due to the fact that the tangential active force alone can drive the ring monomers into the TT motion. Therefore, for smaller $Wi$, t the observed TT is dominated by the active forces. Interestingly, at the intermediate $Wi$, the type-I and type-II exhibit a completely different trend in $f_{tt}$. As discussed in the main text, type-I ring experiences a competition between the active tangential force and the shear force that impacts their TT motion. With the increase in $Wi$, this competition becomes substantial, and $f_{tt}$ starts decreasing until it reaches a minimum. To understand this minimum, please

refer to (Fig. S15(b)). One can see that at $Wi \approx 3700$, the oscillations in $C_{x_i/L}$ are feeble compared to $Wi \approx 750$, indicating the stalling of the TT motion. Such stalling is not observed for type-II activity.

### S5 Movie Description

- **Movie 1** (movie1.mp4): Tumbling motion in type-II active ring at $Wi = 370$ and $Ac\text{-}II = 10$.
- **Movie 2** (movie2.mp4): Pure tank-treading motion of type-I active ring at $Wi = 370$ and $Ac\text{-}I = 10$. For clear visualization of TT motion, a small segment of the ring has been marked in red color.
- **Movie 3** (movie3.mp4): Stalling of type-I active ring at $Wi = 7500$ and $Ac\text{-}I = 10$.
- **Movie 4** (movie4.mp4): Type-II active ring showing a mix of tumbling and tank-treading dynamics at $Wi = 7500$ and $Ac\text{-}II = 10$.
